# Maternal inheritance of primary sex ratios in the dark-winged fungus gnat *Lycoriella ingenua*

**DOI:** 10.1101/2024.11.08.622740

**Authors:** Maria Shlyakonova, Katy M. Monteith, Laura Ross, Robert B. Baird

**Affiliations:** Institute of Ecology and Evolution, University of Edinburgh, Edinburgh, EH9 3TJ

## Abstract

Sex determination mechanisms in insects are extraordinarily diverse, although most species have zygotic genotypic sex determination where sex is established by sex chromosomes upon fertilisation. Dark-winged fungus gnats (Diptera: Sciaridae) are a large and speciose family of flies where sex determination is a result of an unusual interplay of zygotic, maternal, and environmental factors. This causes some species to produce clutches of offspring that deviate considerably from the standard 1:1 Fisherian sex ratio. An early study suggested that these primary sex ratios may be heritable from mother to daughter, but this observation has not been corroborated and the genetic basis for this trait remains unknown. Other studies have found that in some species, there is an additional temperature effect on the primary sex ratio, but again the mechanism is unknown. Here, we perform sibling crosses and temperature-shift experiments in a recently isolated line of the species *Lycoriella ingenua* and find evidence for highly variable and heritable primary sex ratios, but no significant environmental effect. We discuss the consequences of our findings for understanding the mechanisms that produce these unusual sex ratios, and the evolution of sex determination more broadly in this clade.

## Introduction

Sexual reproduction is an ancient feature in eukaryotes, yet the mechanisms by which offspring sex can be determined are strikingly diverse (Bachtrog *et al*., 2014), and this is particularly true of insects (Blackmon *et al*., 2017). Most sex determination systems are genotypic, where loci on the sex chromosomes determine offspring sex. For example, in the housefly *Musca domestica*, a Y-linked factor acts as the primary signal for sex determination (Hediger *et al*., 1998); in the fruit fly *Drosophila melanogaster*, it is the dosage of an X-linked transcript (Erickson & Quintero, 2007). There are, however, many exceptions to the rules. For example, genotypic sex determination (GSD) does not always involve distinct sex chromosomes (Weber & Capel, 2021), a single master switch gene (Moore & Roberts, 2013), or even the evolution of separate sexes (Ghiselin, 1969). In some systems sex is not genotypically determined at all, but rather environmentally determined (environmental sex determination, ESD), such as in many reptiles (Bull, 1980), fish (Godwin & Roberts, 2018), and some crustacea (Kato *et al*., 2011). Such diversity demonstrates the dynamic nature of sex determination systems in nature, which undergo frequent turnover in many clades (Vicoso, 2019). The sex determination mechanism of a system directly influences the primary sex ratio: where sex determination is governed by X and Y chromosomes, the segregation of those chromosomes decides the primary sex ratio (Werren & Beukeboom, 1998). While Fisherian sex ratio theory predicts that frequency-dependent selection should result in a 1:1 primary sex ratios, there are many exceptions, notably in Hymenoptera (wasps, bees and ants) (King, 1987; Meunier *et al*., 2008). Often, ESD is a cause of unorthodox sex ratios (Charnov & Bull, 1989; Korpelainen *et al*., 1997), although ESD is extremely rare in insects.

The dark-winged fungus gnats comprise a family of flies (Diptera: Sciaridae) that seem to contradict virtually all perceived wisdom regarding chromosome inheritance and sex determination. They have a complicated chromosome inheritance system called paternal genome elimination (PGE), a form of haplodiploidy where males inherit, but do not transmit, their paternally-inherited chromosomes (Metz, 1938). While oogenesis is normal, every sperm produced by a male bears two copies of the maternally inherited X and one copy of the maternally-inherited autosomes. After fertilisation, all zygotes therefore have three X chromosomes. Sex is determined post-fertilisation in the early cleavage divisions, when either one or two X chromosomes, which are always derived from the sperm, are eliminated, giving rise to males (X0) or females (XX), respectively (**Figure 1**). The elimination of X chromosomes occurs prior to zygotic genome activation (de Saint Phalle & Sullivan, 1996), so is presumably governed by factors that are maternally-deposited into the embryo.

**Figure 1.**
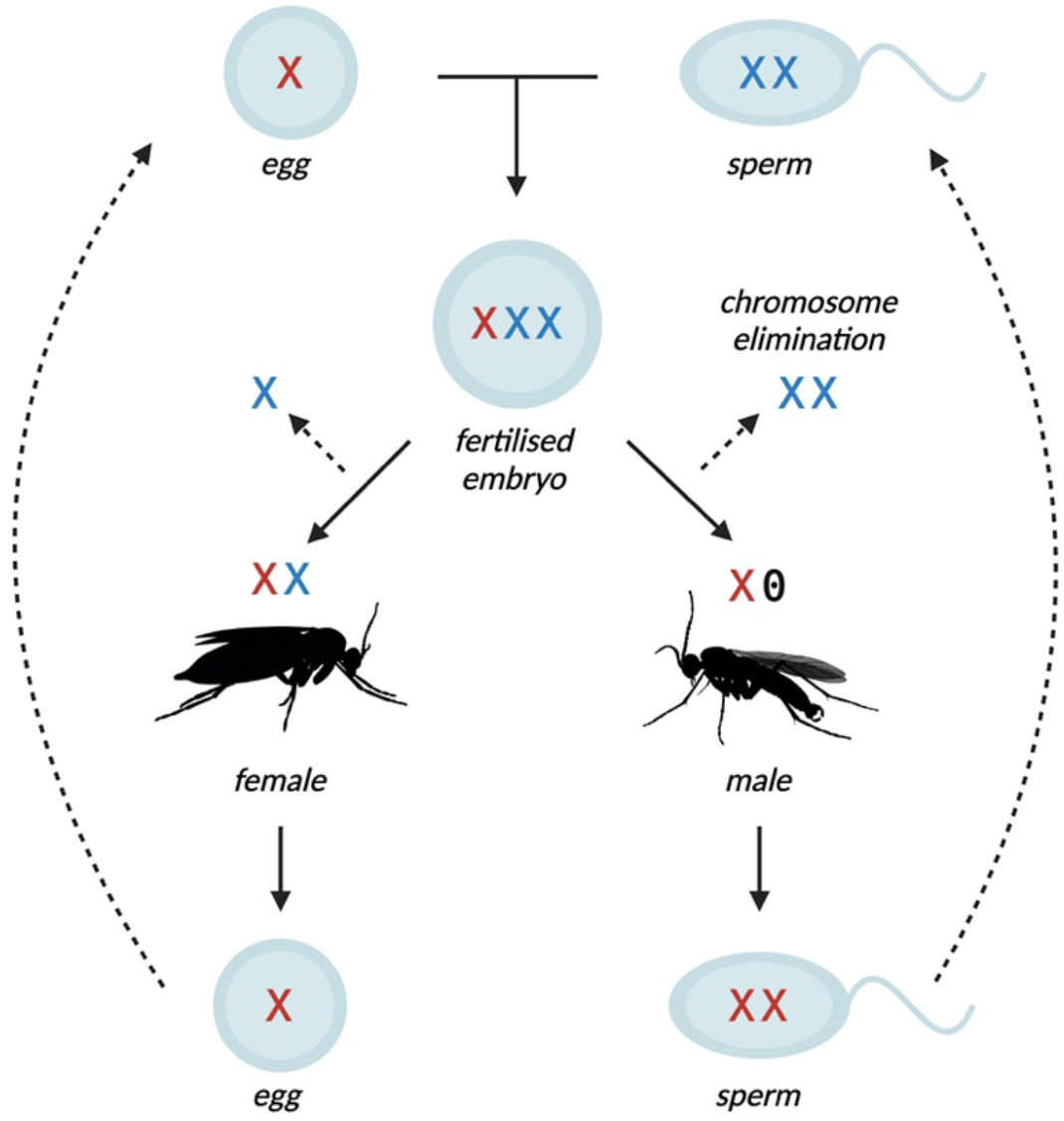
Simplified chromosome cycle of dark-winged fungus gnats (Sciaridae), showing inheritance of the X chromosomes. XXX zygotes form from the fusion of XX sperm and X eggs. Embryos that undergo elimination of one X chromosome develop into females (XX), those that lose two develop into males (X0). The first male meiotic division involves elimination of all paternally-inherited chromosomes; the second division involves nondisjunction of the X. As a result, males always produce sperm with two copies of the X.

As a result, the sex ratio of a brood depends on how many embryos eliminate one or two X chromosomes. In some species, such as *Bradysia coprophila* or *B. impatiens*, mothers produce broods of exclusively one sex (monogenic reproduction). Male-producers and female-producers are genotypically distinct, with the latter harbouring large inversion-based supergenes for which they are always heterozygous (Baird *et al*., 2023b). Other species, such as *B. ocellaris* or *B. reynoldsi*, produce mixed-sex broods (digenic reproduction) (Metz, 1938). In digenic species, sex ratios are known to be highly variable. Single broods that are strongly sex-biased in both directions have been reported in *B. ocellaris* (Davidheiser, 1947; Nigro *et al*., 2007), *B. matogrossensis* (Rocha & Perondini, 2000), and *B. odoriphaga* (Cheng et al., 2017). Monogenic species are also known to produce occasional ‘exceptional’ offspring of the wrong sex (Metz & Schmuck, 1929), suggesting that the distinction between monogeny and digeny may be one of degree, rather than kind. We have previously suggested that these observations of variable sex ratios imply that the proportion of a female’s embryos that develop into males versus females may have an additive genetic component (Baird *et al*., 2023a).

Interestingly, results from a previous study investigating *B. ocellaris* indicated that daughters produce similar sex ratios to their mothers when mated to male siblings (Davidheiser, 1947), suggesting that primary sex ratios may be heritable. However, these experiments have not yet been repeated in any dark-winged fungus gnat species, and heritability of sex ratios in this clade has not been verified statistically. Previous studies have also found a temperature effect on primary sex ratios in the digenic *B. ocellaris* (Nigro *et al*., 2007) and *Lycoriella auripila* (Farsani *et al*., 2013). The temperature-sensitive period was found to be during the late pupal stage when oogenesis is taking place (Berry, 1941) and is the result of conversion of one sex into another rather than sex-biased mortality (Nigro *et al*., 2007). However, the temperature effects appear to be species-specific, with an increase in temperature causing more female-production in *B. ocellaris*, and both high and low temperatures resulting in more males in *L. auripila*.

Dark-winged fungus gnats therefore offer an opportunity to study sex determination where there is a unique interplay of zygotic, maternal, and environmental factors. In the present study, we isolated a line of the digenic species *Lycoriella ingenua* and measured sex ratios in isofemale lines over six generations in order to examine the extent of variability and heritability of primary sex ratios. We also performed temperature-shift experiments to determine whether there is an effect of temperature on sex determination in this species. We find evidence of maternal inheritance of highly variable primary sex ratios in this species, but we do not find a significant temperature effect on the sex ratio. Our results have implications for understanding sex determination in this clade and provide a strong indication that maternal control of offspring sex in sciarids may be under the control of multiple loci.

## Materials and methods

### Insect collection and husbandry

We used a lab culture of *Lycoriella ingenua* (Dufour 1839) that was derived from a wild population collected from Mycobee Mushroom Farm in North Berwick, UK, in February 2022. The species was tentatively identified as *Lycoriella* by inspecting clasper morphology (Broadley *et al*., 2018) and confirmed as *L. ingenua* with a BLASTn (Altschul *et al*., 1990) search of the cytochrome oxidase subunit 1 (COI) barcode sequence (Folmer *et al*., 1994) using WGS data generated for another study (Baird et al. *in prep*). In the literature *L. ingenua* is described as being a digenic species under the synonym *Sciaria mali* (Metz, 1938), which we confirmed after rearing isofemale lines for a generation. The fly stocks are maintained in mass cultures (i.e. several individuals of each sex per vial) at 18°C and 70% relative humidity in 28mm × 95mm polypropylene vials containing 2.2% Bacteriological agar. Larvae are fed a mixture of brewer’s yeast, mushroom powder, spinach powder, and ground straw; a small pinch is added to vials every day 2-3 days during larval development, from the day of egg hatching until pupation. The life cycle (egg to adult) lasts approximately four to five weeks (Gerbi, 2024).

### Primary sex ratios in isofemale lines

We established isofemale lines from mass cultures of *L. ingenua* at random: one male and one female were placed inside a vial and left to mate. We originally set up 47 isofemale G0 lines, of which 42 produced eggs that hatched and survived to produce F1 offspring. Following pupation, vials were checked daily for eclosing F1 adults. We recorded the primary sex ratio as the proportion of male offspring that eclosed from each vial.

We established sibling crosses using eclosed F1 adults from 28 of the 42 successful lines: we aimed to set up 5 sibling crosses per clutch (i.e. 135 total crosses, although not all survived to the next generation). The above process was repeated to generate F2 primary sex ratio counts, and again for F3 and F4 counts, but setting up 3-5 sibling crosses per clutch, depending on the number of available offspring (Figure 2a).

**Figure 2.**
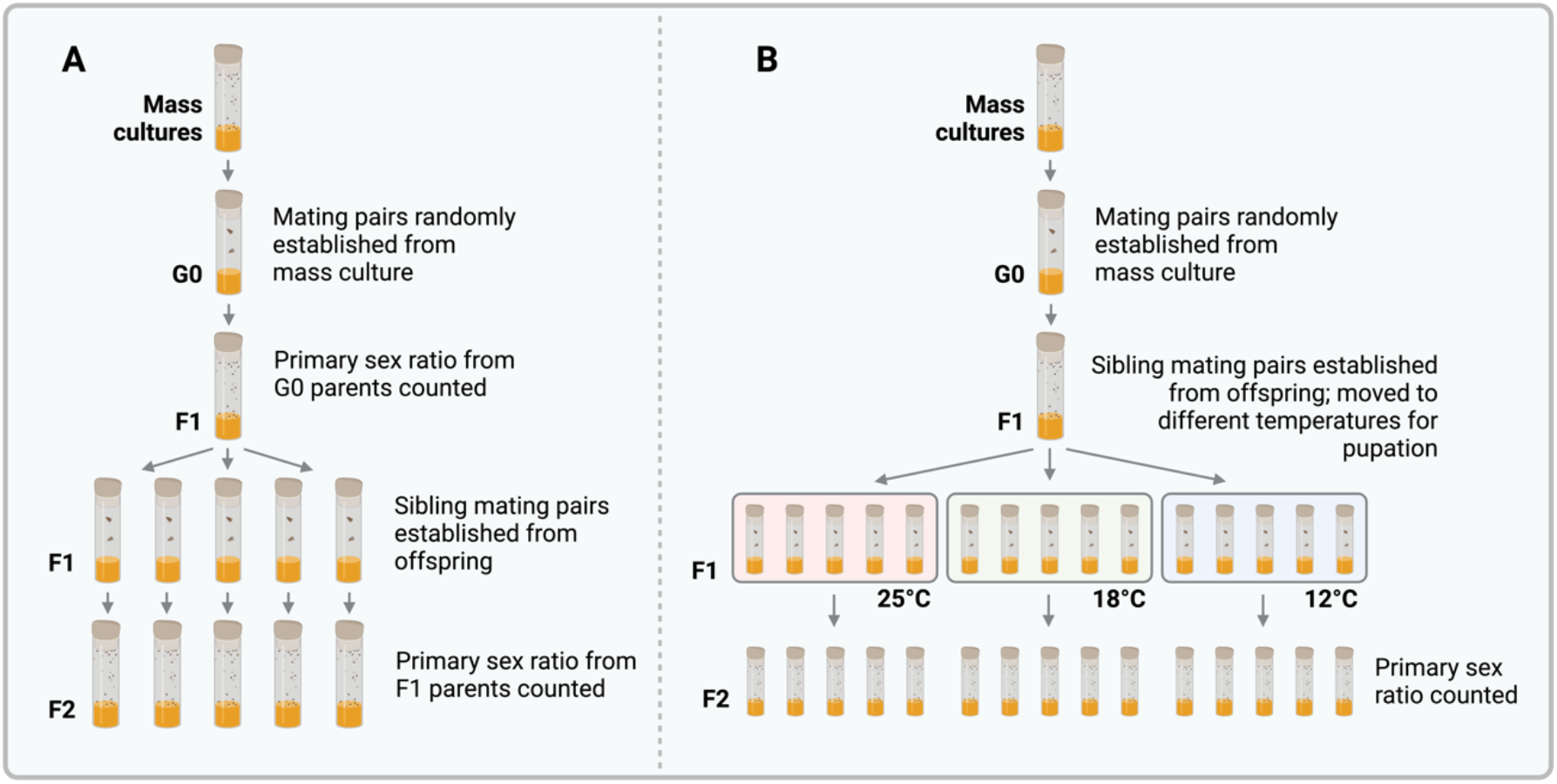
(A) Experimental design to examine heritability of primary sex ratios. After establishing isofemale lines from mass cultures, sibling crosses were set up over successive generations and offspring counted; the last flies to be counted were the F4 generation (i.e. offspring of F3 females). (B) For the temperature experiment flies were either held at 18°C or moved to 25°C or 12°C for the duration of pupation (oogenesis). Upon emergence, females that had been subjected to different temperatures for pupation were mated and their offspring counted.

### Temperature-shift experiments

We also used F1 offspring from the primary sex ratio experiment to perform temperature-shift experiments. We established 15 F1 sibling crosses from each of the 28 lines (total of 405 crosses) and up to 5 crosses from each F1 clutch were randomly allocated to one of three different temperatures; 12°C, 18°C, and 25°C. Previous studies report that temperatures above 27°C are lethal to dark-winged fungus gnats, and that temperatures below 12°C slow development by several months (Reynolds, 1938; Nigro *et al*., 2007). We designed the temperature shifts to encompass the pupal stage of the mother, when oogenesis occurs, which has previously been shown as the stage at which temperature affects the sex ratios (Nigro *et al*., 2007). This was achieved by moving the developing flies from 18°C, the control temperature, to the shift temperature at the onset of pupation. The eclosing females were then mated with male siblings, and their offspring (the F2 generation) were reared at 18°C and sex ratios were recorded upon eclosion.

### Statistical analyses

All data visualisation and analysis was done in RStudio (R Core Team, 2023). To test if primary sex ratios deviated from the Fisherian 1:1 expectation, we performed binomial tests for each brood and combined *P* values using Fisher’s method. To determine whether primary sex ratios were heritable, we tested for a correlation between the sex ratios produced by mothers and their daughters using a linear regression, weighting primary sex ratios by brood sizes. We tested whether siblings produced more similar primary sex ratios than non-siblings by calculating and weighting the absolute differences in sex ratios for siblings and non-siblings and performing a Welch two-sample *t* test on the weighted differences. We further assessed whether the variance of sex ratios within each isofemale line (i.e. all individuals derived from the same founding female) was lower than expected by random chance, with more weight given to larger broods. To test this, we generated 1,000 random sex ratio distributions for each isofemale line under the null hypothesis of random variation, and these simulated distributions were used to create a null distribution of variance. We then calculated the observed variance in the actual sex ratio values for each line, compared this observed variance to the null distribution, and computed *P* values as the proportion of bootstrapped variances that were lower than the observed variance.

We examined changes in sex ratios across generations using a mixed-effects model, with sex ratio as the response variable and generation as a fixed effect, and conducted subsequent pairwise comparisons between generations. To see if brood size (i.e. offspring mortality) changed over generations, we used mixed-effects model with brood size as the response, generation as a fixed effect, and founder line as a random effect, and then conducted pairwise contrasts between successive generations. We used a generalised linear model to determine if the decline in brood size over generations was sex-biased, with brood size as the response variable, an interaction between generation and sex as a fixed effect, and founder line as a random effect.

To determine whether there was an effect of temperature on the sex ratio we used a linear regression model with primary sex ratios weighted by brood size. We also checked for changes in mortality in different temperature treatments using a linear model, and also for changes in sex-specific mortality using a linear model with offspring sex as an interaction term.

## Results

### Variability and changes in primary sex ratios and survivorship

We found that sex ratios were significantly more variable than expected under a normal binomial distribution (Fisher’s combined *P* > 10^−135^). Although the mean and median proportion of male offspring were across all progenies were 54% and 52%, respectively, primary sex ratios varied from 0% to 100% male offspring (**Figure 3A**). Interestingly, between the founder generation (G0) and their offspring (F1), broods became significantly more male-biased (*P* < 0.0001), but between generations F1 and F3, they became significantly more female-biased (F1-F2 comparison: *P* < 0.001; F2-F3 comparison: *P* < 0.0001) and significantly less male-biased (both comparison: *P* < 0.001; **Figure 3B**). We found that overall mortality increased over the first two generations (*P* < 0.0001), which could result from inbreeding. Since inbreeding could affect male and female fitness differently (Ebel & Phillips, 2016), these changes in primary sex ratios over generations could be due to sex differences in offspring mortality. However, we found that the extent of decline in offspring count over generations did significantly differ between males and females, suggesting this is not the case. We also found an increase in brood sizes between generations F3 and F4 (*P* < 0.05), which could represent recovery after purging of genetic load and loss of lethal allele combinations (Mongue *et al*., 2016; **Figure 3C**).

**Figure 3.**
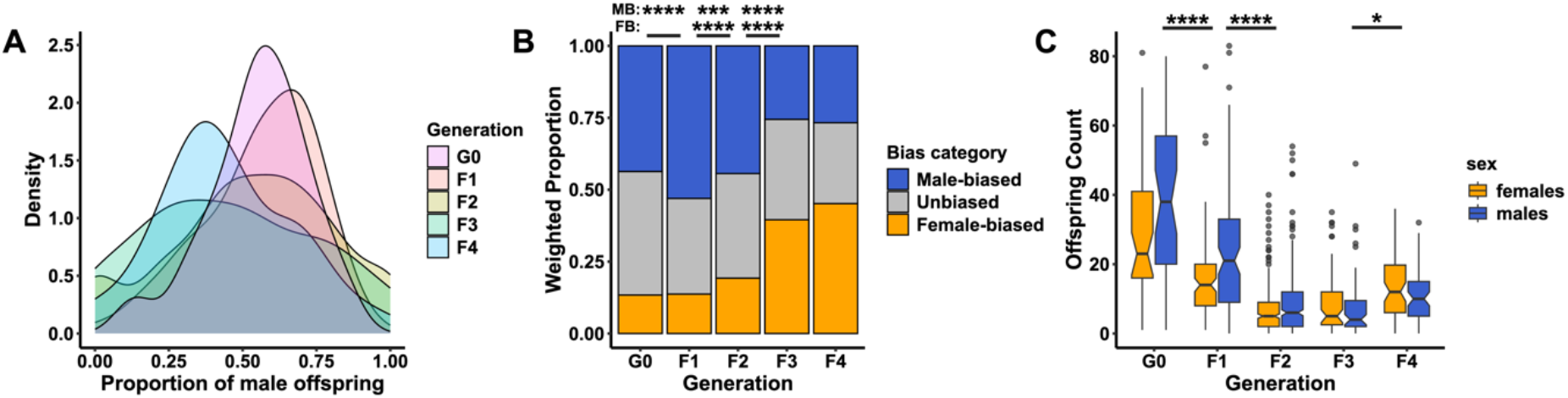
Sex ratio variance and survivorship. **A** Density plot showing variation in primary sex ratios for the five generations studied. **B** change in sex ratio bias over generations (weighted by brood size), with asterisks showing comparisons between male-(MB) and female-biased (FB) clutches between generations. **C** Male and female offspring counts over generations, with asterisks showing statistical comparisons of clutch size between successive generations. Asterisks represent significance levels (\*\*\*\**P* < 0.0001; \*\*\**P* < 0.001; \**P* < 0.05).

### Inheritance of the primary sex ratio

Females that were mated to their male siblings tended to produce similar primary sex ratios to their mothers. specifically, we found a statistically significant correlation between the primary sex ratio produced by a mother and that of her daughter (*P* < 10^−10^), suggesting that primary sex ratios are heritable in *L. ingenua* (**Figure 4**). In further support of this, primary sex ratios were significantly more similar between siblings compared to between non-siblings (Welch two-sample *t* = -9.5, *df* = 1892.5, *P* < 10^−15^), and in 18 of the 28 isofemale lines, primary sex ratios were more significantly more similar to one another than expected by chance (**Supplementary Figure 1**).

**Figure 4.**
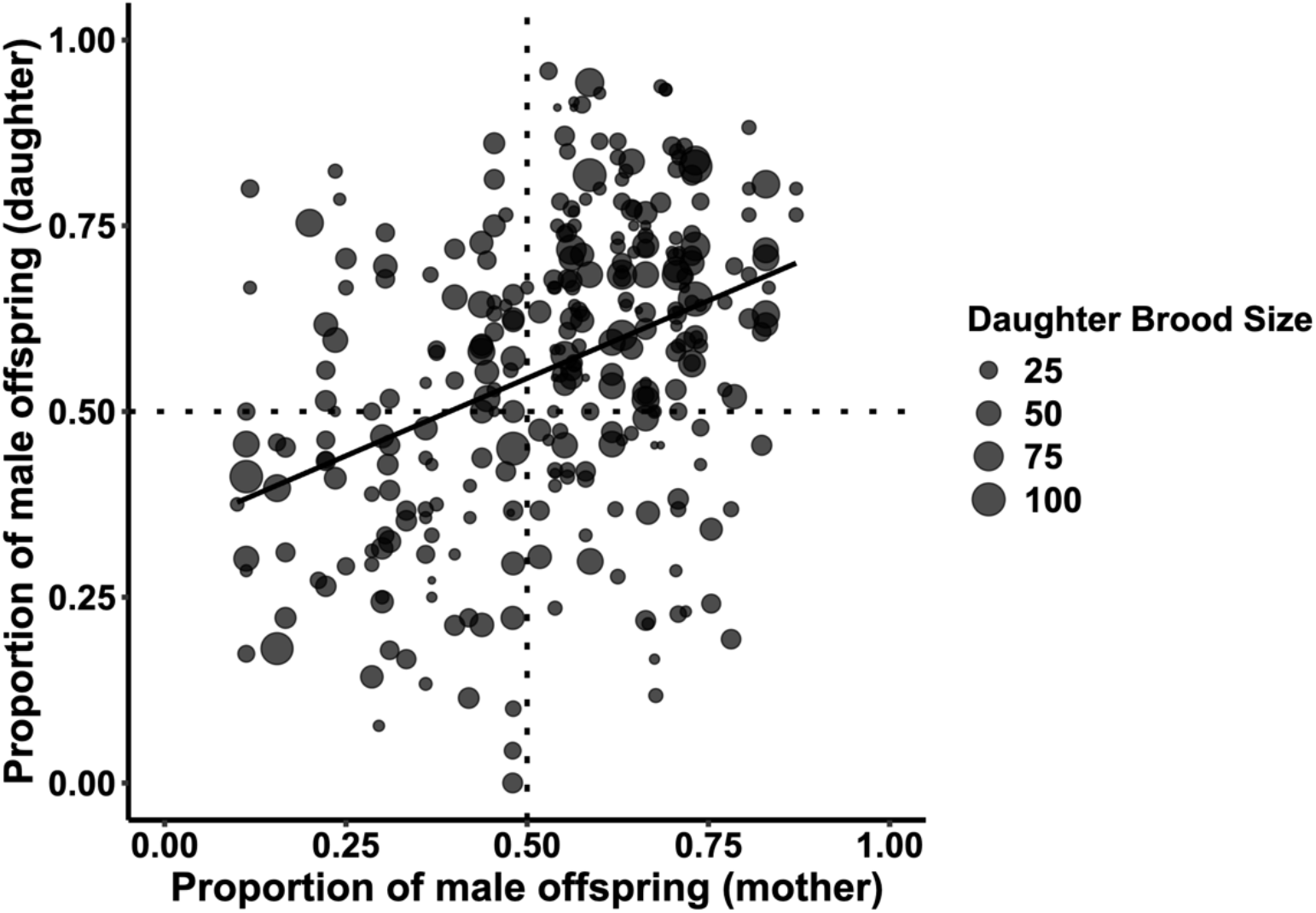
Mother versus daughter primary sex ratios (only brood sizes >= 10 are shown). Each point represents the proportion of male offspring in a mother-daughter pair. The regression line was plotted using the formula = ‘y ∼ x’.

### Temperature effects

The median proportion of males produced by mothers reared at 12°C, 18°C, and 25°C was 47%, 56%, and 50%, respectively. We did not find a significant effect of temperature on primary sex ratios (*P* = 0.971, **Figure 5A**). We also tested for differences in mortality under the different temperature treatments, and found that offspring mortality was lower at both 12°C and 25°C compared to 18°C, but the difference was only significant for the 12°C-18°C comparison (**Figure 5B**). There was no significant difference when considering only female offspring mortality, but there was for males at 12°C, suggesting that males are perhaps more sensitive to changes in the temperature at which the mother was reared (**Figure 5C**).

**Figure 5.**
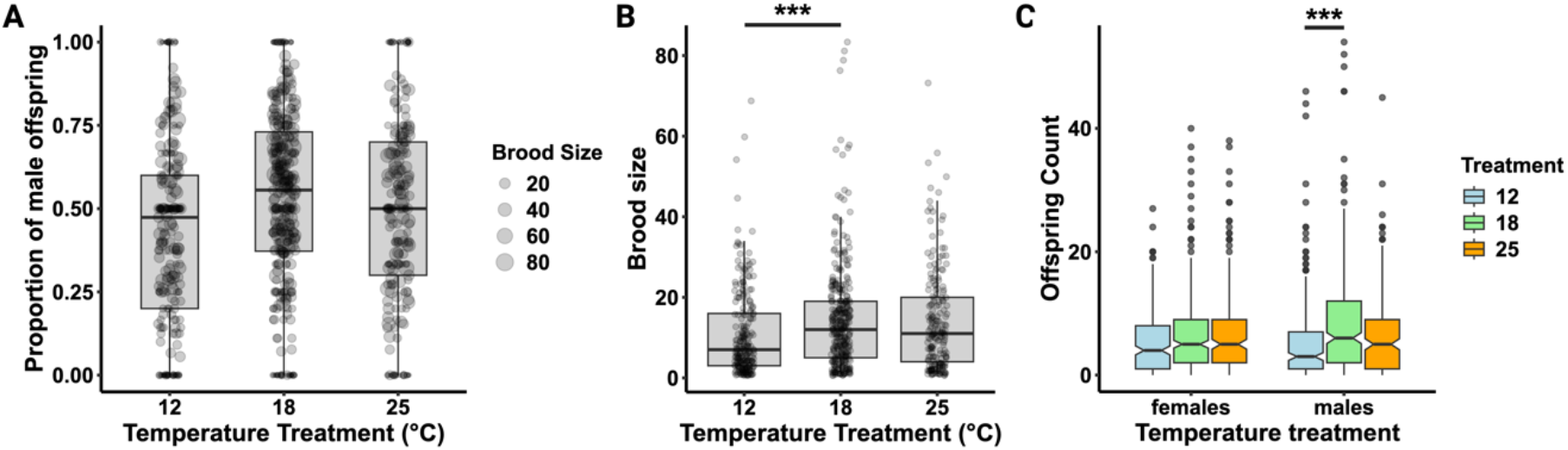
Temperature effects on development. **A** Effect of temperature on primary sex ratios. Dotted lines mark 50:50 sex ratios. **B** Effect of temperature on brood size. **C** Effect of temperature on male and female counts.

## Discussion

Dark-winged fungus gnats have an unusual system of sex determination where a combination of zygotic, maternal, and environmental factors influence the sex of offspring (Sánchez, 2010). Sperm with two X copies fertilise eggs with one X, producing XXX zygotes that proceed to lose either one or two X chromosomes in early embryogenesis, developing as XX females or X0 males, respectively. Whether a zygote develops as male or female can be influenced by the maternal genotype (Baird *et al*., 2023b) and temperature during oogenesis (Nigro *et al*., 2007; Farsani *et al*., 2013). In digenic sciarids, where females produce mixed-sex broods, mothers are known to produce variable sex ratios (Rocha & Perondini, 2000; Nigro *et al*., 2007; Farsani *et al*., 2013), and results from a previous study suggest that the primary sex ratio may be heritable (Davidheiser, 1947). In the present study we sought to determine the extent of variability, heritability, and the effect of temperature on primary sex ratios in a recently cultured laboratory strain of the species *Lycoriella ingenua* with the aim of improving our understanding of the genetic and environmental mechanisms that determine sex in this unusual clade of flies.

We found that primary sex ratios are highly variable in this species, ranging from 0% to 100% male offspring. This is concordant with observations from studies of other sciarid species, including *B. ocellaris* (Nigro *et al*., 2007), *B. odoriphaga* (Cheng *et al*., 2017), *B. matogrossensis* (Rocha & Perondini, 2000), *L. agraria* (Farsani *et al*., 2013), and *Scatopsciara cunicularis* (Sawangproh & Cronberg, 2016). That the primary sex ratio of a brood appears to be a continuous trait suggests that it is under the control of more than one locus, as suggested by Rocha & Perondini (2000) and expanded upon in Baird *et al*. (2023a). Moreover, our finding that primary sex ratios are more similar between siblings compared to between non-siblings, and that daughter sex ratios correlate significantly with mother sex ratios, afRirms that determination of the primary sex ratio has a heritable genetic component. However, there was considerable variation between siblings in primary sex ratios, suggesting that perhaps many loci or indeed environmental factors may be at play.

Some of the primary sex ratios we observed were particularly extreme, and we found some evidence that sex ratios became more skewed over successive generations. In other digenic sciarids, progenies of only one sex are frequently reported (Davidheiser, 1947; Rocha & Perondini, 2000; Nigro *et al*., 2007). Some other sciarid species, in contrast, produce strictly single-sex broods (Metz, 1938; Lara *et al*., 1965; Steffan, 1974), and in two species this is known to be associated with large chromosomal inversions (Carson, 1946; Crouse, 1979). In Baird *et al*. (2023a), we proposed that these inversions evolve to ‘trap’ alleles that influence the sex ratio, resulting in a transition from digenic to monogenic reproduction, and that this may be a result of sex ratio selection. Documented cases of multi-locus or polygenic sex determination are rare and are thought to be only a transient phenomenon (Schartl *et al*., 2023; although see Kocher *et al*., 2024). Polygenic sex determination is thought to be inherently unstable (Rice, 1986; Bateman & Anholt, 2017), which is exacerbated when environmental effects such as temperature influence offspring sex (Van Dooren & Leimar, 2003). In Sciaridae, the evolution of monogeny from digeny may be one way or resolving this instability.

In contrast to some previous studies on other sciarid species (Nigro *et al*., 2007; Farsani *et al*., 2013), we did not find a significant of effect of temperature on primary sex ratios, suggesting that this temperature effect may be species-specific. The precise mechanism of temperature effects on sciarid sex determination is yet to be explored. Maternal factors that determine the sex of an embryo must be transferred from nurse cell cytoplasm to oocytes. Extreme temperatures may affect the rate of transfer as suggested by Nigro *et al*. (2007) and Sánchez (2010), but to our knowledge the effect of temperature on maternal mRNA or protein deposition has not been studied. An alternative possibility is that extreme temperatures induce meiotic nondisjunction. Errors in female meiosis may result in eggs with aberrant karyotypes: since all eggs fuse with an XX sperm, nullisomic eggs would produce XX zygotes and disomic eggs XXXX zygotes. In typical systems this would be fatal. However, under this system of sex determination, post-zygotic X elimination would restore some eggs to a viable ploidy (e.g. if one X is eliminated from XX zygotes or if two Xs are eliminated from XXXX zygotes). Rearing *Drosophila melanogaster* at low or high temperatures has been known to induce meiotic nondisjunction (Grell, 1979), mostly affecting the X chromosome (Tokunaga, 1970). Furthermore, exceptional male and female offspring that occur in the monogenic species *B. coprophila* can result from sex-chromosome nondisjunction during oogenesis, producing aneuploid eggs (Crouse, 1960).

### Conclusion

Inheritance of primary sex ratios is a rare phenomenon in nature, since in most organisms females produce standard Fisherian 1:1 sex ratios. Paternal transmission of primary sex ratios has previously been reported in haplodiploid Hymenoptera (Werren & Stouthamer, 2003) and has been observed outside of insects in copepods (Voordouw *et al*., 2005), polychaete worms (Premoli *et al*., 1996), and fairy shrimp (Beladjal *et al*., 2002). In *Formica* fire ants, supergenes are known to underlie split sex ratios (production of either future queens or drones) (Lagunas-Robles *et al*., 2021). Here we present, to our knowledge, the first study of maternal inheritance of primary sex ratios in a diploid organism. We confirm that primary sex ratios in this unusual sex determination system are variable, have a heritable genetic basis, and are likely determined by multiple loci, but we do not find a significant temperature effect. Future work should focus on understanding the loci involved and the mechanism by which sex determination occurs in sciarids. More generally, this system offers excellent opportunities to explore polygenic control of sex determination, as well as the interplay between zygotic, maternal and environmental contributions to sex determination.

## Acknowledgements

We would like to thank Bea Herliczka at Mycobee Mushroom Farm for providing access to collect fly samples. We thank Andrew Mongue for useful comments on the draft manuscript, as well as members of the Ross Lab for discussions and for help in maintaining fly stocks. All authors were funded by an ERC starting grant (PGErepro, to LR).

## Author contributions

RBB collected and established *Lycoriella ingenua* cultures. MS, RBB and LR conceived of the study and experimental design. MS and KMM collected data. RBB performed analysis and wrote the draft manuscript. All authors reviewed the final manuscript.

## Data availability

Data produced in this study and code used in analyses are available in the following GitHub repository: https://github.com/RossLab/Lycoriella_sex_ratios.

