## Supplementary Figure 1 for "Maternal inheritance of primary sex ratios in the dark-winged fungus gnat *Lycoriella ingenua*"

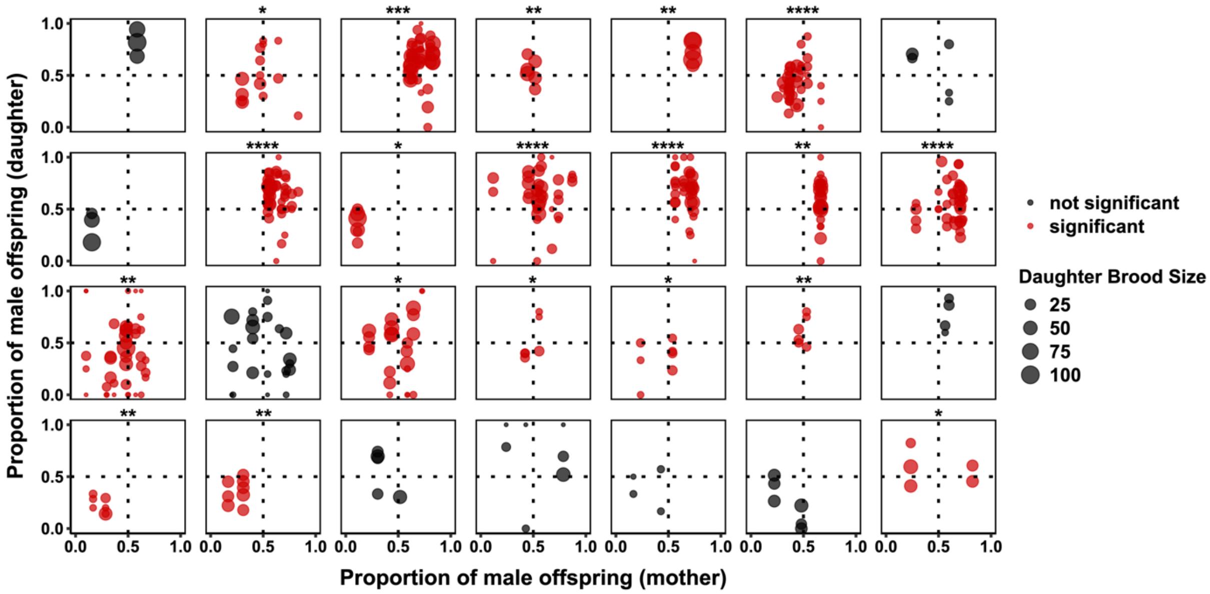


**Supplementary Figure 1.** Mother versus daughter primary sex ratios, separated by isofemale line (i.e. all broods within a plot are derived from the same founding female). Each isofemale line is coloured based on whether the variance of sex ratios within a line are significantly more similar to each other than expected by random chance, with more weight given to larger brood sizes. Asterisks represent significance levels (*$P$ < 0.05, **$P$ < 0.01, ***$P$ < 0.001, *$P$ < 0.0001).
